# Dissecting the cortical stages of invariant word recognition

**DOI:** 10.1101/2025.03.28.645903

**Authors:** Aakash Agrawal, Stanislas Dehaene

## Abstract

Fluent reading requires the brain to precisely encode the positions of letters within words, distinguishing for instance *FORM* and *FROM* across variations in size, position, and font. While early visual areas are known to encode retinotopic positions, how these representations transform into invariant neural codes remains unclear. Building on a computational model of reading, we used 7T fMRI and MEG to reveal a cortical hierarchy in which early visual areas (V1–V4) predominantly encode retinotopic information, whereas higher-level regions, including the Visual Word Form Area (VWFA), transition to an ordinal letter-position code. MEG analyses confirm that retinotopic encoding emerges early (60–200 ms), followed by a shift toward ordinal representations in later time windows (220–450 ms). Despite this transition, word position remained a dominant factor across all time points, suggesting a concurrent coding of both retinotopic and abstract positional information. These findings uncover the spatiotemporal dynamics by which the human brain transforms visual input into structured linguistic representations, shedding light on the cortical stages of reading and their developmental and clinical implications.

## Introduction

Fluent reading requires extensive practice that harnesses and refines the brain’s visual circuitry for linguistic purposes (1). The acquisition of reading leads to the emergence of a specialized region along the ventral visual pathway, often referred to as the Visual Word Form Area (VWFA), that exhibits selective responses to the statistical regularities of the learned script(s) (2–5). Indeed, imaging studies show that, once literacy has been mastered, the VWFA responds robustly to written words and sublexical features (2, 5–7), and does so invariantly across major transformations in size, spacing, font and, partially, retinal position (8–12).

A central puzzle, however, is how the human brain transitions from retinotopic representations - where letter identity is tightly coupled to a particular location on the retina - to more abstract, ordinal representations that preserve letter order despite shifts in the visual field (13, 14). Neuroimaging research indicates that early visual areas (V1–V4) faithfully encode retinal coordinates (15). More anterior regions such as the VWFA, however, exhibit only minimal sensitivity to screen position, implying that their letter-position codes are more “word-centered” (8, 13, 16, 17). This progression parallels the broader hierarchy of areas for object recognition along the ventral stream, wherein regions progress from encoding localized low-level features to constructing position-invariant, high-level representations (18, 19). Despite widespread acceptance that the ventral pathway supports increasingly sophisticated encoding, the precise stage at which letter-position codes decouple from retinotopy and the degree to which retinotopic signals might linger at higher levels remain active areas of inquiry (10).

Recent computational modelling has shed additional light on the neuronal circuits that may underlie the transformation from retinotopic to abstract invariant representations. In our prior work (20), we simulated the acquisition of a VWFA-like selectivity by training convolutional neural networks (CNNs) to classify written words, thus implementing the recycling of an illiterate visual system in a literate one (21, 22) ((see Fig 1A). In the later layers of the CNNs, we discovered two broad classes of units: those that were selective to specific letters irrespective of position, and those coding for letter positions in an ordinal manner, with sensitivity to an approximate ordinal position relative to either the beginning or the end of the word (23). This shift from predominantly retinotopic to ordinal-based coding supports longstanding cognitive models of reading that emphasize the importance of order-based codes, with “slots” for successive letters regardless of their actual size and position on screen (24, 25). Nonetheless, we also uncovered residual position dependence in later layers (20), consonant with imaging findings that even the VWFA may encode coarse positional cues (10).

**Figure 1:**
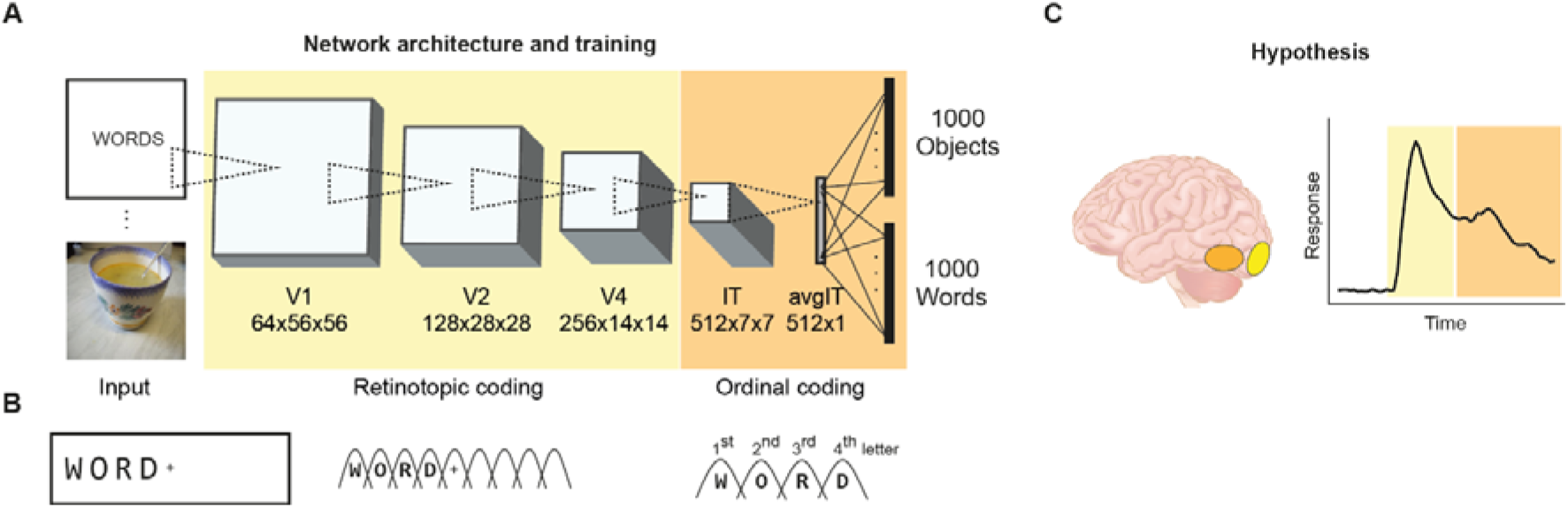
Hypothetical mechanism for invariant word recognition in brains and machines. (A) To simulate literacy acquisition, the CORnet-Z convolutional neural network (CNN) architecture, initially used to simulate image recognition, was trained to recognize both written words and pictures (20, 23). Compared to an “illiterate” network trained to recognize only pictures, this literate network acquired units tuned to letters and words, particularly in its deeper layers (IT and avgIT). (B) Schema of the retinotopic and ordinal coding schemes discovered in this CNN. From the given input “WORD”, the initial stages process letters at their retinotopic position; only downstream areas rely on an abstract ordinal code which combines letter identity with ordinal position relative to word endings. (C) Based on those neural network simulations, we hypothesize that early regions of the ventral visual pathway, in an early time window, primarily encode retinotopic information. In contrast, downstream regions, in a later time window, transition to encoding ordinal relationships.

From a psychophysical perspective, recent evidence also suggests a compositional neural code for reading, whereby the representations of words can be partly predicted by combining abstract letter identities (26–28). Reading acquisition increases the separability of letter codes for consecutive letter, thus making strings like AB and BA increasingly different for the trained reader (26). Orthographic codes that preserve the relative order of letters appear crucial to distinguishing near-anagrams and handling the multitude of typographic changes encountered in daily reading (29). Yet direct neuroimaging evidence pinpointing where and when retinotopic encoding yields to ordinal (or more abstract) coding has been limited.

In the present study, we build on computational predictions (20) to examine precisely how these letter-position codes evolve in vivo in the human brain. By combining ultra–high-field (7T) functional MRI and time-resolved magnetoencephalography (MEG), we disentangle the contributions of absolute letter location, whole-word position and ordinal letter position in shaping cortical responses to short letter strings. By using specialized stimuli that manipulate retinotopic and ordinal positions systematically, we test the prediction, derived from our computational modelling, that lower-level areas maintain highly position-specific representations while higher-tier regions converge on ordinal letter sequences anchored by edge letters (13, 20, 21, 27).

Our findings demonstrate a gradual shift in letter-position coding along the ventral visual pathway, as revealed by our combined use of 7T fMRI and MEG. Early visual areas (e.g., V1–V4) retained strong retinotopic signals tied to a letter’s precise location, whereas more anterior regions, including the VWFA, exhibited an emerging sensitivity to the order of letters within words. MEG analyses further showed that retinotopic representations emerge early (within ∼60 ms of stimulus onset), while ordinal coding becomes more prominent in later time windows. By matching these neuroimaging results against CNN models trained to read, we provide direct evidence that the human brain gradually transitions from a retina-bound encoding of letter identity toward a word-centered representation that still preserves position information but emphasizes the ordinal sequence of letters. This account thus clarifies how visual cortex balances invariance and precision, enabling effortless recognition of words regardless of peripheral changes.

## Results

### Computational models

In our previous work (20, 23), we trained a variant of the CORnet-Z convolutional neural network (CNN) to recognize both objects and written words. This training procedure led to the emergence of word-selective units broadly analogous to those observed in the human ventral visual pathway. Here, to deepen those findings, we probed these units with carefully designed letter strings ((see Figure 2A) aimed at dissociating two broad classes of position-coding schemes: a *retinotopic* scheme, in which the position of each letter is anchored to its location on the retina, and an *ordinal* scheme, in which each letter’s position is encoded relative to the beginning and/or end of the string. We restricted stimulus selection to three letter pairs comprising a frequent letter and another rare dissimilar one in French (OX, TB, and EQ), i.e. the very same stimuli used in brain imaging, but similar results were observed with other letters.

**Figure 2:**
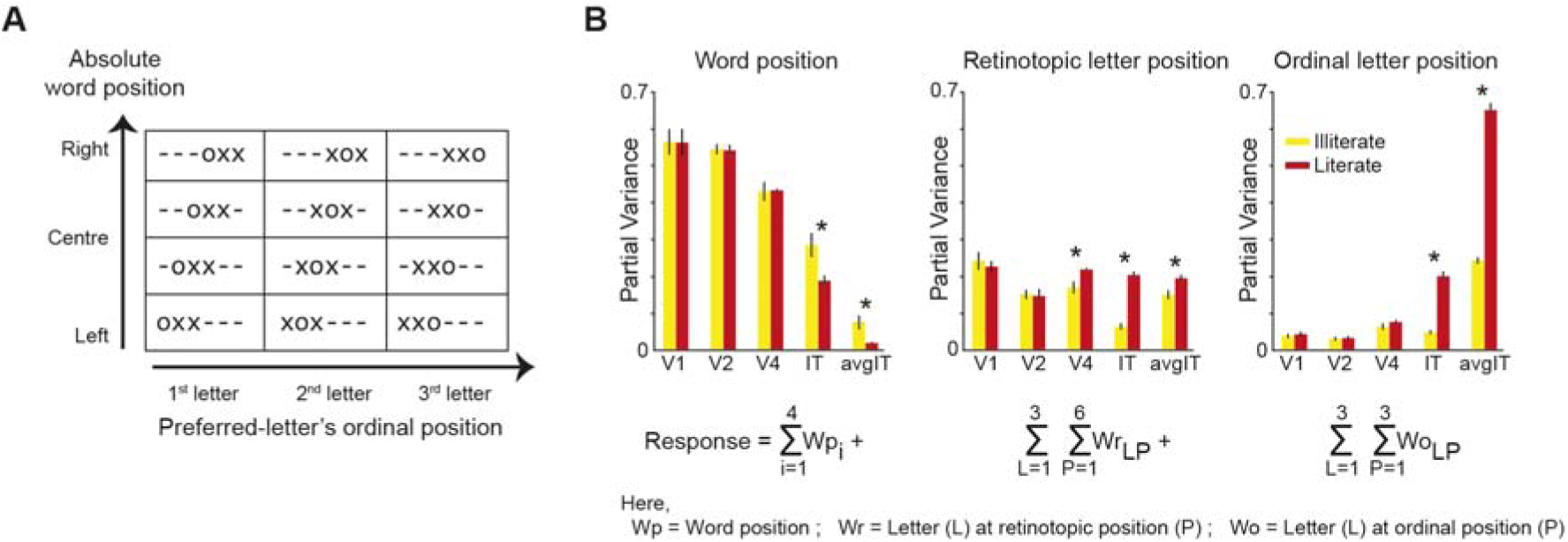
Specific stimuli dissociate position-coding schemes in CNN models of literacy acquisition. (A) Schematic of the stimuli used to dissociate different position-coding schemes. We present 3-letter strings consisting in a frequent letter (here the letter 0) and two infrequent letters (here X). Across 12 different stimuli, we vary the absolute string position (rows) and the ordinal position of the preferred letter (columns). In addition to the illustrated stimuli, two other letter triplets (EQQ and TBB) were used, resulting in a total of 36 unique letter string stimuli. The ‘-’ symbol represents a blank space and was not shown on screen. (B) Modelling the responses of literate and illiterate CNNs to those stimuli. Bars show the proportion of variance explained, averaged across word-selective units, for different factors in the encoding model. Literate CNNs, red bar; illiterate, yellow bars. The linear model predicts the response of each unit as a combination of absolute word position, retinotopic letter position, and ordinal letter position. As shown in (A), the model includes four unique word positions, six retinotopic positions, three ordinal positions, and three unique letter identities (O, E, T). Error bars denote the standard deviation across five network instances, and asterisks (*) indicate a significant difference between literate and illiterate networks. The model shows that absolute string position and retinotopic letter position dominate in early layers, while the deep layers IT and avgIT are based on ordinal coding. With literacy training, the word position code decreases while the letter X retinotopic position and especially the letter X ordinal position codes increase.

Our prior results showed that illiterate networks (trained only on objects) relied almost exclusively on retinotopic coding, whereas literate networks (jointly trained on objects and words) gradually shifted toward ordinal coding in deeper layers. Notably, however, some units in both networks exhibited mixed selectivity and did not neatly fall into either category based on simple tuning profiles alone. To address these mixed responses more rigorously, we developed a linear encoding model incorporating three main factors - word position, retinotopic position, and ordinal position - together with letter identity (see Methods). For each word selective unit, we measure the neural response to each of the 36 stimuli (3 letter identities X 3 ordinal position X 4 word positions) and fitted it with a hierarchy of models, using LASSO optimization to account for potential collinearity. To quantify the unique contribution of each factor, we first measured the variance explained by word-position terms alone. We then added retinotopic-position terms and computed the change in explained variance relative to the baseline model. Finally, we included ordinal-position terms to estimate their unique explanatory power. Note that the model was fit only once, and variance decomposition was performed hierarchically to isolate the contribution of each factor. Because illiterate networks lack word-selective units, we restricted this analysis to the units identified as word-selective in the literate condition and tracked their responses across network layers. Akin to localization of VWFA, a unit is identified to be word selective if its responses to words were greater than the responses to other image categories (likes faces, objects, tools, etc.) by at least three standard deviations.

Consistent with our previous observations, the *early layers* of the literate network were dominated by word-position factors, while *ordinal coding* emerged strongly only in the deeper layers. The magnitude of these effects diverged from the IT layer after literacy training. Interestingly, retinotopic terms remained consistent across layers in both literate and illiterate models. We speculate that these retinotopic effects in literate network is amplified (starting from V4 layer) by emergent letter-selective units, which remain partially anchored to visual location. Thus, our linear model not only validated earlier findings of a retinotopic-to-ordinal shift, but also provided quantitative estimates of the unique variance contributed by each factor ((see Figure 2B).

### Letter position coding in the human brain

We next asked whether the progression from retinotopic to ordinal letter-position coding, as predicted by our computational model, could also be observed in the brains of expert readers. To this end, we measured both functional MRI (fMRI) and magnetoencephalography (MEG) signals while participants viewed the same stimuli used to probe our literate and illiterate networks (see Methods). The stimuli were presented in random order, and participants were instructed to maintain fixation on a centrally located red dot and press a button on the rare occasions where all letters were identical (e.g., XXX, BBB, or QQQ; 10% of trials). Subjects performed this task accurately (94.2% in fMRI; 81.1% in MEG) and detected the target strings more efficiently when they appeared near the fovea (Figure S2).

From whole-brain 7T fMRI data (n = 16; 1.2 mm isotropic), using a combination of functional localizer and anatomical maps, we extracted four individual regions of interest (ROIs) along the ventral visual pathway: V1-V3, V4, LOC, and VWFA (see methods). An example subject’s ROIs is shown in (see Figure 3A. Within each voxel of each participant, we applied the same linear encoding model described for the literate network ((see Figure 2), estimating partial variance explained by word position, retinotopic position, and ordinal position in a hierarchical regression framework. To control for overfitting, we estimated model fits after randomly shuffling the response vector, refitted the model, and subtracted the resulting variance explained from the original model fit. This noise estimation procedure was repeated five times (see Methods). As expected, the fraction of variance explained by word position decreased steadily from early visual areas toward VWFA, while the opposite trend was observed for retinotopic and ordinal coding.

**Figure 3:**
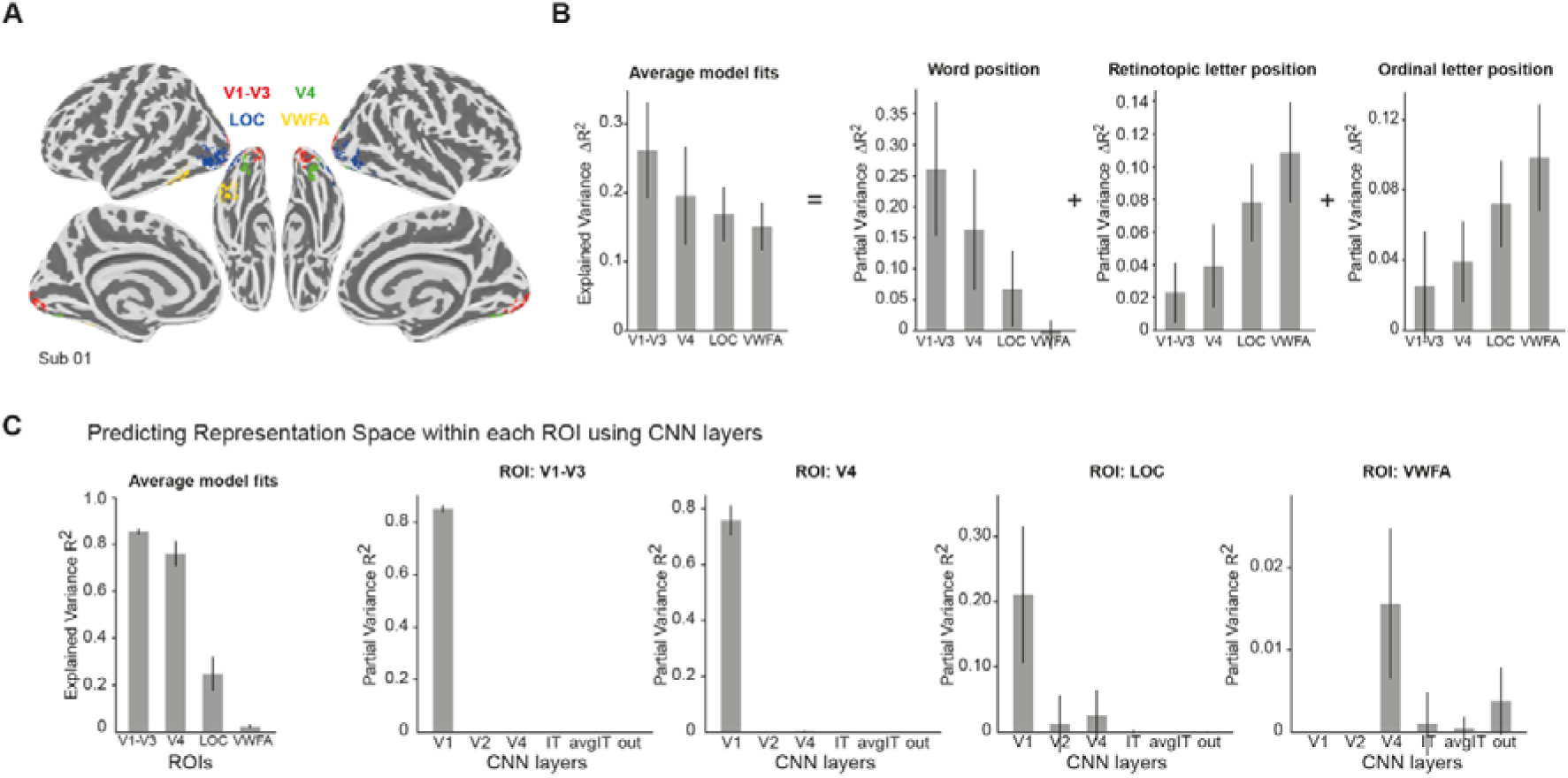
Position coding schemes detected in the human brain using 7T fMRI. (A) Regions of interest (ROIs) for an example subject. Each ROI is functionally defined based on contrasts between experimental conditions and further constrained using anatomical masks. (B) Variance explained by letter x position encoding model, averaged across units with significant model fits (p < 0.01). The decomposition of the variance components suggests a decline in word position coding along the ventral visual pathway, while retinotopic and ordinal letter position coding gradually increase up to the VWFA. (C) Predicting the representation space of stimuli within each ROI using a convolutional neural network (CNN). The left panel shows overall model fits (R²) for each ROI, while the right panels depict the variance explained by different CNN layers within each ROI. Early visual areas (up to V4) are best explained by early CNN layers (V1), whereas the visual word form area (VWFA) is primarily predicted by deeper network layers. Error bars indicate standard deviation derived from 100 bootstrap estimates across subjects.

To further compare the neuroimaging data to our computational model, we computed representational dissimilarity matrices (RDMs) for each ROI and performed a hierarchical regression to determine how well each CNN layer’s RDM predicted the observed neural patterns. Consistent with earlier studies using objects (30), the representation in early visual areas were best predicted by the early layers of the network, and the representation in VWFA best correlated with the later layers of the network, with a peak at V4 layer (Figure S3B), suggesting a gradual transition from retinotopic to ordinal encoding in both the model and human ventral visual pathway.

Using MEG, we tracked how retinotopic and ordinal letter-position coding unfold over time in the human brain. Participants viewed the same letter-string stimuli from the fMRI experiment while we recorded neural responses from −100 to +500 ms relative to stimulus onset. We applied a hierarchical regression model to the MEG signal, averaged in 10 ms bins for each sensor and participant, to capture the distinct contributions of word, retinotopic, and ordinal position factors. The partial variance for each factor was computed relative to a pre-stimulus baseline.

The word position factor became significant first, at approximately 60 ms post-stimulus, followed by the retinotopic position factor around 100 ms. The ordinal position factor emerged significantly later, at about 220 ms, suggesting a sequential progression from coarse positional information to more abstract, context-dependent coding ((see Figure 4A). Guided by these onset times, we divided the MEG epoch into two temporal windows: an early window (60–200 ms) and a late window (220–500 ms). Within each window, we performed a representational similarity analysis analogous to the approach used in the fMRI study (see (see Figure 3B), once again applying hierarchical regression to determine how well each layer of the literate CNN explained the observed neural patterns.

**Figure 4:**
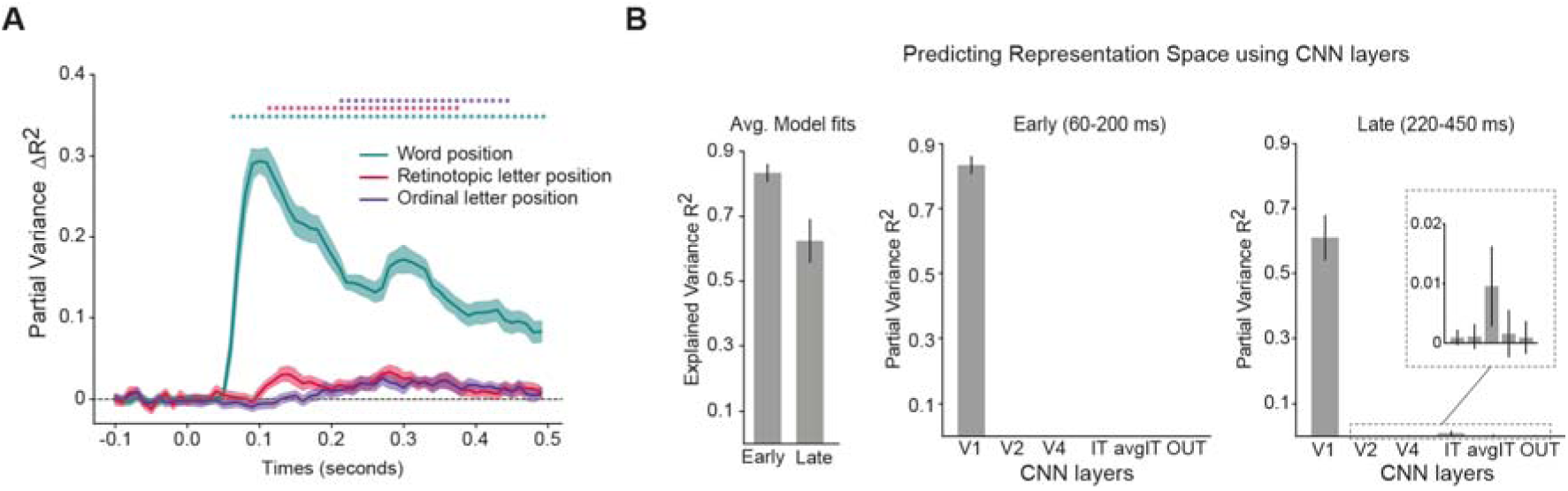
Temporal dynamics of position coding schemes, as measured by MEG. (A) Temporal evolution of encoding model fits from MEG recordings, showing baseline-corrected explained variance (R²) from -100 to 500 ms relative to stimulus onset. Three distinct position encoding schemes are compared: word position (green), retinotopic letter position (red), and ordinal letter position (purple). The explained variance for each scheme was calculated by subtracting prestimulus baseline R values. Shaded areas represent standard error across subjects, with asterisks indicating statistically significant clusters (cluster-based permutation test, p < 0.05). (B) CNN layer-specific contributions to neural representation across temporal windows. The explained variance (R) is shown separately for early (60-200 ms) and late (220-450 ms) processing stages (left). The model’s primary visual layer (V1) accounted for the majority of explained variance in both time windows. The inset highlights the emergence of higher-order visual processing, showing significant contribution from IT layer specifically during the late time window (220-450 ms). Error bars indicate standard deviation derived from 100 bootstrap estimates across subjects.

Our findings indicate that the early window (60–200 ms) was best accounted for by the V1 layer of the network, reflecting strongly retinotopic encoding. While the retinotopic component remained dominant during the late window (220–500 ms) as well, a small yet significant contribution from the IT layer emerged, consistent with the delayed onset of ordinal coding ((see Figure 4B).

In sum, these results demonstrate that although retinotopic coding persists throughout the processing window, the emergence of ordinal coding in later stages suggests an additional abstraction mechanism at work. Note that the partial variance reported here is averaged across all sensors, which substantially reduces the apparent effect size. However, analyzing the variance explained at individual sensors revealed robust effect sizes for each of the three factors: the highest R² was 0.90 for word position, 0.28 for retinotopic position, and 0.24 for ordinal position. Furthermore, topographical maps at peak time points showed that word position is encoded in early visual areas, while ordinal letter positions are encoded in more anterior regions at later time points (Figure S4).

### Searchlight analysis

To integrate spatially localized fMRI activity with the millisecond resolution of MEG, we conducted a searchlight analysis linking local fMRI activation patterns to time-resolved MEG patterns (31). Each voxel within the brain was analyzed along with its surrounding neighborhood of 125 voxels (5×5×5) to construct a representational dissimilarity matrix (RDM) using the correlation distance metric. We then applied hierarchical regression to predict this RDM as a linear combination of RDMs estimated from early (60–200 ms) and late (220–450 ms) MEG responses. The underlying hypothesis was that shared retinotopic variance across MEG time points would be predominantly captured in the early time window, while the late response map would provide an unbiased estimate of ordinal position coding.

A large proportion of variance in early visual areas was indeed predictable using the early time window of MEG, confirming the retinotopic nature of initial cortical representations. Interestingly, the late time window showed dominant contributions in the left hemisphere, extending up to VWFA, as well as lateral regions in the right hemisphere. These findings reinforce our earlier conclusions that VWFA and deeper regions of the ventral visual pathway encode abstract ordinal position.

To further generalize these findings, we conducted a whole-brain searchlight analysis to systematically explore the neural correlates of CNN layers across the entire cortex. The results revealed a strong correspondence between early CNN layers and early visual areas, supporting their role in initial retinotopic processing. In contrast, later layers of the literate network exhibited increased similarity to higher-order cortical regions implicated in language and reading (Figure S5). This pattern suggests a gradual functional shift from low-level visual processing to increasingly abstract representations that are critical for fluent reading.

**Figure 5:**
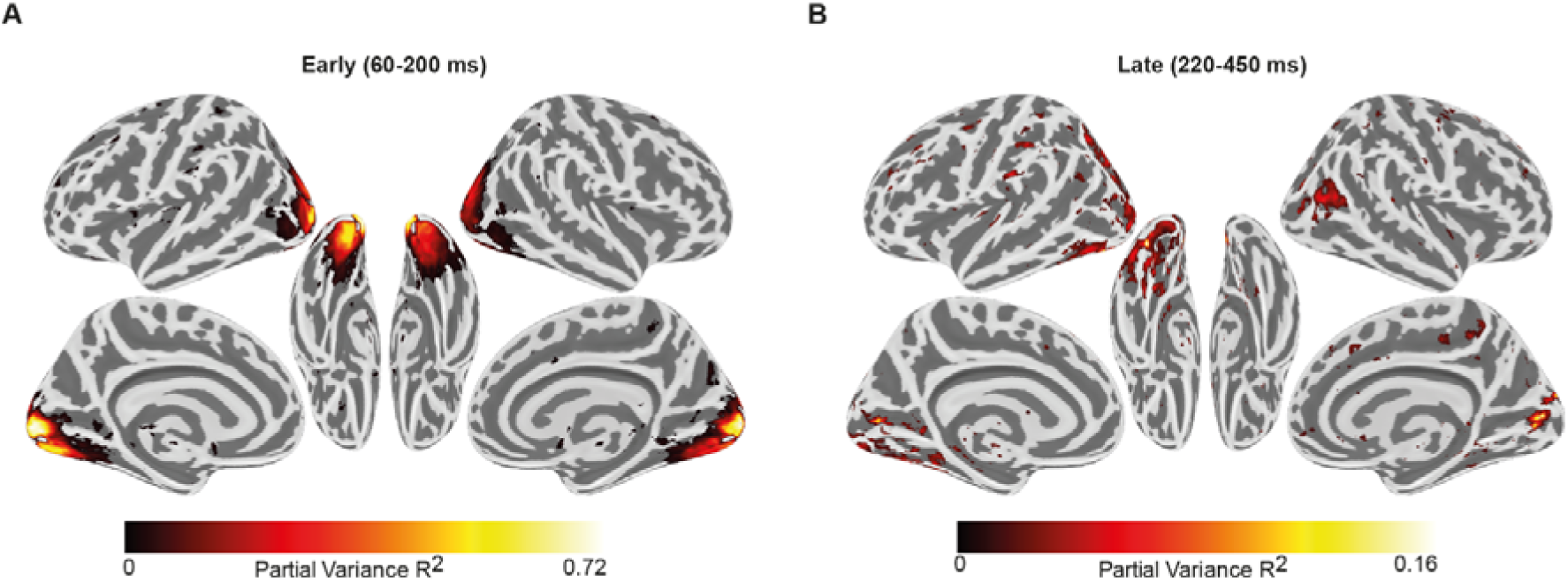
Searchlight analysis between fMRI and MEG. (A) Searchlight analysis revealing spatial correspondence between fMRI and MEG representational patterns. For each searchlight cube of voxels, the local fMRI representation space was modelled as a linear combination of MEG representation space from early and late time windows. The partial variance (R) map from early time window (60-200ms) are thresholded to indicate the significant partial variance after FDR correction (p<0.05) (B) Same as (A) but for late (220-450 ms) time window.

### Decoding analysis

We also examined if our findings could be confirmed using multivariate decoding methods ((see Figure 6). Encoding models, representational similarity and multivariate decoding are complementary methods, each with their own advantages and drawbacks (32, 33). Here, decoding largely confirmed the above findings, yet with reduced sensitivity possibly due to the superimposition of massive retinotopic signals.

**Figure 6:**
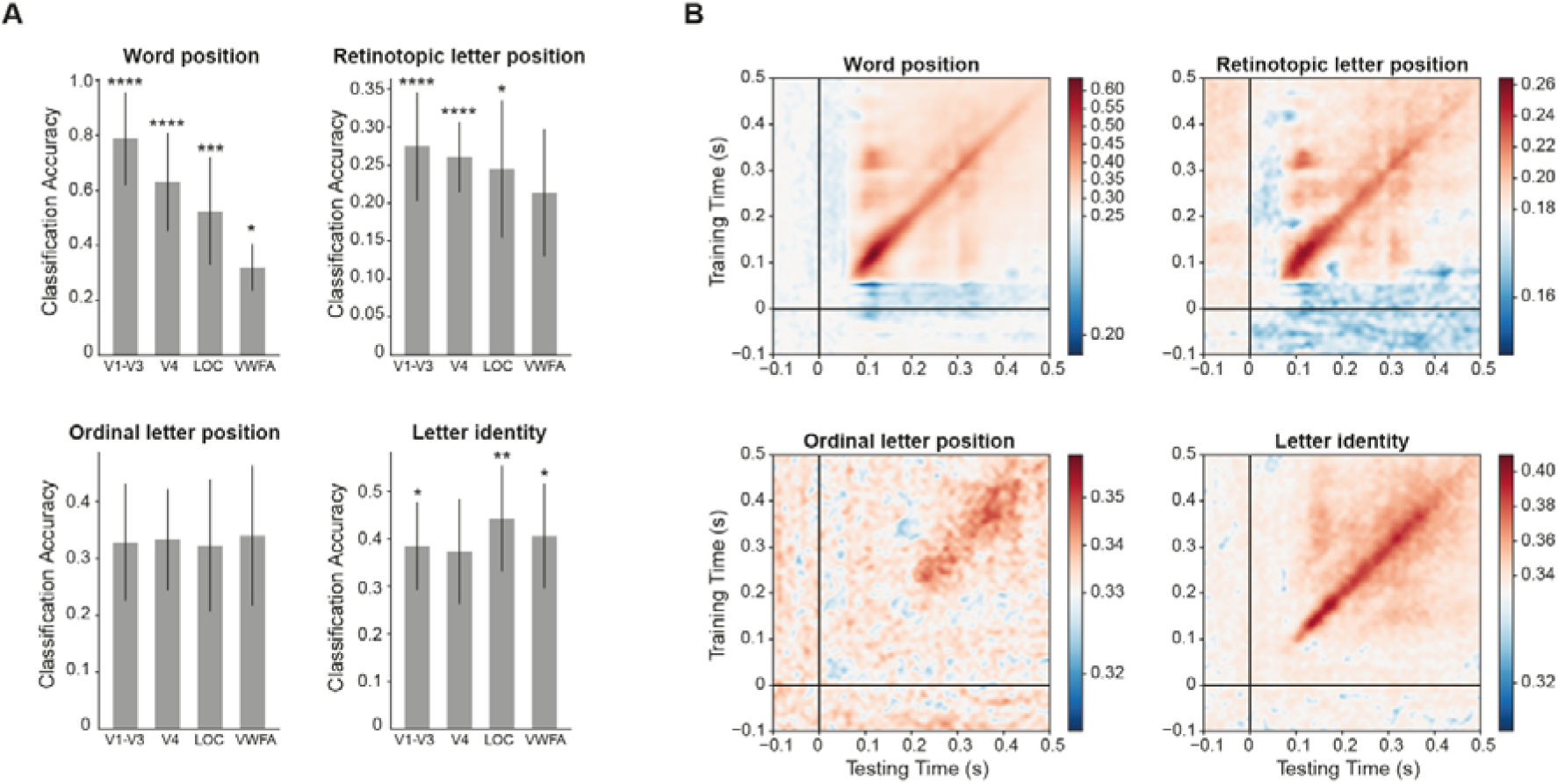
Decoding stimulus properties across space and time. (A) Classification accuracy across different ROIs from fMRI for absolute word position (chance = 25%), absolute letter position (chance = 16.67%), letter identity (chance = 33.33%), and ordinal letter position (chance = 33.33%). Error bars indicate standard error of mean across subjects and the asterisks indicate statistical significance (* p < 0.05, ** p < 0.005, *** p < 0.0005, etc. using one sample t-test). (B) Decoding of the same four codes in MEG. The generalization-across-time matrices (35) indicate the performance of the decoders for different training (y axis) and testing time points (x axis).

In fMRI, within each region, using voxels as features, we fitted a linear classifier to decode the following factors: word position, letter identity, absolute letter position, and ordinal letter position. To avoid overfitting, we performed 3-fold cross-validation (see Methods). Consistent with earlier findings, decoding accuracy for word position was highest in early visual areas, but progressively decreased along the ventral visual pathway, ultimately failing to reach statistical significance within VWFA ((see Figure 6A). The decoding of ordinal letter position was not significant in any regions. The other two factors could be significantly decoded in most of the ROIs, with higher accuracy for letter identity in object and word selective regions than early visual areas (p < 0.005; paired t-test).

MEG signals, again grouped into bins of 10ms each, were also submitted to the same decoding and representational similarity analyses. Decoding accuracy was above chance for word position, absolute letter position, and letter identity starting at ∼60ms and peaking around ∼140ms. However, decoding of ordinal letter position became significant only at ∼220ms, which aligns with the timings of word selectivity in the anterior region of the fusiform gyrus (34). Generalization-across-time matrices (35) exhibited a largely diagonal form ((see Figure 6), indicating that those codes likely propagated to successive brain regions over time. However, while word position and retinotopic letter position decayed over time, both the letter identity and ordinal letter position codes were more sustained, as expected if they are to guide subsequent word identification and processing. Overall, decoding findings confirm the transition from early time windows driven by retinotopy to a later ordinal letter position code.

## Discussion

It is generally accepted that invariant representations emerge as information travels along the ventral visual pathway. While this phenomenon has been well characterized for object categories, particularly faces (36, 37), its specific underpinnings for written words remain to be fully elucidated. In this study, we tested computational hypotheses derived from earlier simulation work (20, 23). Using high-field (7T) fMRI and time-resolved MEG in expert adult readers, we mapped the spatial and temporal transformations that give rise to position-invariant word representations. Consistent with established theories of reading (21), we found that early visual areas (V1–V4) predominantly encode retinotopic information, whereas higher-tier regions such as the left fusiform gyrus transition to more word-centered or ordinal representations ((see Figure 5). Notably, word position remained a robust factor influencing neural activity at multiple time points, suggesting a mixed-selectivity principle (38) where the persistence of position-dependent representations coexist with the emergence of neurons that are attuned to letter order and relative sequence (7, 10, 34).

At both the single-voxel (fMRI) or single-channel (MEG) level, as well as in aggregate population responses, we observed a gradual reduction in word-position sensitivity alongside an increasing emphasis on ordinal coding in more anterior regions of the ventral stream. Although direct decoding failed to reveal significant ordinal codes in the VWFA, encoding models and representational similarity analyses showed that this region’s response patterns aligned closely with later layers of convolutional neural networks explicitly trained for invariant word recognition and which comprise neurons tuned to letter-order. In MEG, retinotopic signals emerged around 60 ms, whereas ordinal coding became evident around 220 ms, and both effects were corroborated by decoding and hierarchical regression. A searchlight analysis confirmed that the contribution of ordinal position in later time windows was specifically localized to the left ventral pathway leading toward the VWFA. While the ordinal effect sizes might appear small in R² terms, their correlation-based equivalents are commensurate with typical findings in fMRI (39).

These results support the empirical validity of our prior simulation work on “literate” convolutional networks trained to recognize both written words and images, which revealed a continuum rather than a sharp transition from retinotopic to ordinal position coding (20, 23). Related outcomes in unsupervised networks (40) further underscore the notion that a combination of edge-letter tuning and letter-based representation underlies efficient visual word recognition (27). Moreover, our searchlight approach showed that early network layers best matched the responses of early visual cortex encoding precise letter locations, whereas intermediate layers corresponded to lateral occipital regions, and the network’s output layer mapped onto a broader language-related reading network, including precentral gyrus (41, 42).

Despite these insights, we acknowledge that the search for fine-grained neural population codes in high-level visual areas remains at the edge of detectability using non-invasive methods, including the state-of-the-art 7T fMRI and MEG techniques used here. Thus, the present work await confirmation using intracranial recordings, which could unambiguously establish the proposed neural coding schemes. Ultimately, only single-unit recordings could provide a specific test of the precise neural mechanism proposed to lie at the origin of ordinal coding, i.e. the existence of intermediate neurons sensitive to “space bigrams”, i.e. units that detect the presence of a specific letter at an approximately fixed distance relative to the space that marks the beginning or the end of a written word. Future research should examine mid-tier regions of ventral visual pathway to determine how precisely such space-based coding might contribute to the emergence of ordinal representations.

Although our study focused on short, controlled stimuli, we anticipate that the same neural mechanisms would extend to more ecologically valid reading of longer words or continuous text. In fact, this might lead to improved effect size as real words typically elicit stronger activations and engage additional language-related circuits compared to the orthographically implausible strings that we were forced to use here in order to isolate the location of a single frequent letter (2, 5, 6, 34). Likewise, while the CNNs in our simulations contained no recurrent layers, the delayed emergence of ordinal coding (∼220 ms) is likely to reflect, at least in part, reverberating and top-down influences onto the reading circuitry. Nonetheless, the timing and localization of these signals concur with previous fMRI (13), MEG (43) and intracranial recordings (7, 34) studies of word processing.

An important future direction will be to elucidate how these position-coding schemes mature in developing readers or degrade in reading impairments. Several subtypes of dyslexia are associated with the orthographic level in multiple languages (44–48). In letter-position dyslexia, children make errors that swap nearby letters, particularly inside words, for instance reading “form” as “from” (45, 47, 49–51). These errors resemble the letter-transposition errors that also occur in normal readers under time pressure or subliminal priming (12, 52–55). They can be easily explained, at an orthographic coding level, by an imprecision in the proposed invariant letter-position code: with increased variance of the proposed ordinal tuning curves, inner letters would be allocated imprecise locations, while the first and last letters would remain sufficiently sharply encoded. A second type of reading difficulty is attentional dyslexia, which occurs in both acquired and developmental forms (44, 47, 56). In this case, letter position *within* a word is highly preserved, but the errors swap letters between two nearby words (e.g. “let saw” is misread “set law”). Again, this is similar to the illusory conjunction errors that occur in normal readers under time pressure (57). In the present framework, such errors might be explained by an inappropriate attentional selection of the word to be read and inhibition of the surrounding words. If their neural codes were mixed, there would be more than one candidate letter for each position, explaining why, in this specific reading impairment, letters migrate across words while maintaining their within-word positions. Thus, several error types may arise from a partial breakdown in the shift from retinotopic to more abstract positional codes. In the future, cross-linguistic comparisons with scripts of varying structure, transparency and linearity (5, 26, 58, 59) may clarify how the writing system’s visual structure modifies the balance between location-based and location-invariant responses.

Taken together, our findings provide a detailed understanding of how letters are anchored in retinotopic space at early latencies and progressively recast into word-centered, ordinal representations in higher regions of the ventral visual pathway. By uniting predictions from literacy-trained neural networks with precise neuroimaging data, we may ultimately arrive at a biologically grounded account of how letter-position encoding supports fluent reading, distinguishing even minimal anagrams while tolerating variations in font, spacing, and size. More broadly, our work illustrates how the cortex’s general-purpose machinery for visual recognition can be recycled for orthographic processing (1, 4, 22), capitalizing on early retinotopy while leveraging higher-order abstraction. This framework sharpens our understanding of typical reading, illuminates the neural origins of reading disorders, and underscores how cultural inventions like written language harness and reshape the inherent plasticity of the human brain.

## Methods

### Model architecture and training

Among the many available convolutional neural networks that can predict neural responses along the ventral visual pathway, we chose CORnet-Z architecture for two reasons: 1) It has a modular structure that resembles the stages of processing in the visual cortex (V1, V2, V4, IT, avgpool IT, output), 2) It has fewer parameters, thus lowering the training time while achieving high levels of accuracy on synthetic word datasets (23). Similar to our previous work, we first trained this network on the ImageNet dataset (phase 1), which contains ∼1.3 million images across 1000 categories. This was considered an illiterate network, which encodes the visual properties of objects but not text. Next, we extended the number of output nodes to 2000 (1000 images + 1000 words), with full connectivity to H layer units, and retrained the entire network jointly on ImageNet and a synthetic word dataset (phase 2), which also contained 1.3 million images of 1000 words. This was considered a literate network. To estimate the variability across training sessions, the literate network was trained starting from the same five instances of illiterate networks. Pytorch libraries were used to train these networks with stochastic gradient descent on a categorical cross-entropy loss. The learning rate (initial value = 0.01) was scheduled to decrease linearly with a step size of 10 and a default gamma value of 0.1. Phase 1 training lasted for 50 epochs and phase 2 training for another 30 epochs. The classification accuracy did not improve further with more epochs.

### Stimuli

To improve network performance, ImageNet images were transformed using standard operations such as “RandomResizedCrop” and “Normalize”. The images were of dimension 224×224×3. To avoid cropping out some letters in the word dataset, the default scale parameter of RandomResizedCrop was changed such that 90% of the original image was retained. For fair comparisons, other operations such as flipping were not performed on the Imagenet dataset as the same operation would create mirror words in the Word dataset, which is not typical in reading.

The French words included frequent words of length between 3-8 letters. The synthetic dataset comprised 1300 stimuli per word for training and 50 stimuli per word for testing. These variants were created by varying position (-50 to +50 along the horizontal axis, and -30 to +30 along the vertical axis), size (30 to 70 pts), fonts, and case. A total of 5 different fonts were chosen: two for the train set, i.e. Arial and Times New Roman and three fonts for the test set, i.e. Comic Sans, Courier, and Calibri.

### Identification of word selective units

Similar to fMRI localizer analysis, a unit was identified as word selective if its responses to words were greater than the responses to nonword categories: faces, houses, bodies, and tools by 3 standard deviations. The body and house images were taken from the ImageNet dataset. For tools, we used the “ALET” tools dataset, and Face images were taken from the “Caltech Faces 1999” dataset. We randomly chose 400-word stimuli and 100 images each from the other categories for identifying category-selective units.

### Dissimilarity measure

For each layer, we vectorized the activation values, and estimated the pair-wise dissimilarity value using correlation metric i.e., d = 1-r. where r is the correlation coefficient between any two activation vectors.

### Linear model

The stimuli consisted in three-letter strings that shifted across four positions along the horizontal axis, creating four distinct word positions. Each three-letter string therefore spanned up to six potential letter locations on the image, and included three possible ordinal positions (left, center, right). We selected three sets of letter triplets (OX, TB, EQ), with each set comprising one frequent letter and one infrequent letter, yielding a total of 36 unique stimuli.

This design led to a total of 31 predictors in the linear model: 4 word-position factors, 6 retinotopic-position factors × 3 letters, and 3 ordinal-position factors × 3 letters. Formally, for each three-letter stimulus, the observed response vector y (36×1) was modeled as **y=X**β, where **X** is a 36×31 design matrix, and β is a 31×1 vector of unknown parameters. By fitting the full model with LASSO and applying hierarchical regression, we isolated the unique variance explained by word position, retinotopic position, and ordinal position.

In fMRI, estimating model accuracy can be confounded by overfitting, particularly when the number of predictors is large. To account for this, we conducted a noise-based control analysis by shuffling the response matrix y across trials, thereby breaking the true correspondence between stimuli and neural responses. We then refitted the model with the shuffled data and computed the variance explained. This shuffled variance estimate, reflecting the model’s fit to noise, was subtracted from the variance explained by the original (unshuffled) model to obtain a corrected measure of explained variance. To ensure robustness, we repeated this procedure five times with different random shuffles and averaged the results.

### fMRI data

#### Participants

A total of 16 adult subjects (aged b/w 18-40 years) who were native French speakers participated in this study. They were right-handed and had normal or corrected-to-normal vision with no history of any psychiatric disorders or reading difficulties. Participants provided written informed consent for the fMRI study and received monetary compensation. The study was approved by the local ethics committee in the NeuroSpin Centre (CPP 100055) and was conducted following the Declaration of Helsinki.

#### Stimuli

The functional localizer block consisted of various stimuli, including French words, objects, scrambled words, and scrambled objects. For each run, 12 unique images were randomly chosen from a pool of available images. The scrambled images were created by scrambling the phase of the Fourier-transformed images and then reconstructing them using the inverse Fourier transform.

The event block comprised 36 unique stimuli, which consisted of three-letter strings (e.g., OXX, XOX) with specific characteristics. Each string contained one frequent letter (e.g. O) and two repeated rare letters (e.g. X). We specifically selected three letter pairs for this purpose: OX, TB, and EQ, with the first letter in each pair being frequent and the second being rare in French, the language of our participants. These letter pairs were also chosen based on their visual dissimilarity and low bigram frequency. The design was founded upon the hypothesis that the frequent letter would cause activation in putative letter-selective units, while variations in its absolute location on the screen and in its relative location within the string would allow us to establish the relative or positional nature of its neural code. A 3×4 design of the letter strings was used to dissociate the absolute and ordinal positions of the frequent letters (Figure S1). Absolute word position was varied by shifting the strings by one letter at a time, ranging from completely left to completely right of the fixation dot (four levels of word position, resulting in six levels of absolute letter position). At each word position, the odd letter could appear in each of three positions within the 3-letter string (left, center, or right).

The stimuli were presented on a BOLD screen (Cambridge Research Systems, Rochester, UK), a 32-inch MRI-compatible LCD screen with a resolution of 1920 × 1080 pixels. The screen had a refresh rate of 120 Hz and was positioned at the head-end of the scanner bore. Participants viewed the stimuli through a mirror attached to the head coil.

#### Task design

The localizer run comprised 14-s mini blocks in which a total of 14 images were presented for 0.8 s with an inter-stimulus interval of 0.2 s. There were 12 unique stimuli and 2 of them repeated at random time points, corresponding to targets for the one-back task. Following each block of stimuli, a blank screen with a fixation cross was presented for 6 seconds. Each block (words, objects, and their corresponding scrambled versions) was repeated three times within each run. There were two runs of the localizer.

In the event-related runs, each of the 36 stimuli was displayed for 0.2 seconds, followed by a blank screen lasting 2.8 seconds. Within each run, each stimulus was repeated five times. To actively engage the participants, twenty task trials were also included, during which subjects were required to respond with a button press when all the letters in a stimulus were identical (e.g., XXX, BBB, QQQ). Additionally, to introduce variability in the inter-stimulus interval, 10% of the trials (n = 20) did not present any stimulus. This helped to jitter the timing between stimuli. There was a total of 4 runs, 660 s long, and each run started and ended with a blank screen displaying a fixation cross for 4s. The high-resolution MP2RAGE anatomical images were obtained in the middle of the scan session i.e., after two event-related and one localizer run. The subjects were instructed to close their eyes and relax during the anatomical scan.

#### Data acquisition

The Brain images were acquired using a 7-T Magnetom scanner (Siemens, Erlangen, Germany) with an a1Tx/32Rx head coil (Nova Medical, Wilmington, USA) at the NeuroSpin Centre of the French Alternative Energies and Atomic Energy Commission. Dielectric pads were placed around the ear to reduce the signal dropout around the anterior Ventral Occipital Temporal cortex.

Functional data were acquired with a two-dimensional (2D) gradient-echo echo-planar imaging (EPI) sequence using the following parameters: repetition time (TR) = 2000 ms, echo time (TE) = 21ms, voxel size = 1.2 mm isotropic, multiband acceleration factor = 2; encoding direction: anterior to posterior, iPAT = 3, flip angle = 75, partial Fourier = 6/8, bandwidth = 1488 Hz per pixel, echo spacing = 0.78 ms, number of slices = 70, no gap, reference scan mode: GRE, MB Leak Block kernel: off, fat suppression enabled. To correct for EPI distortion, a five-volume functional run with the same parameters except for the opposite phase encoding direction (posterior to anterior) was acquired immediately before each task run. Participants were instructed not to move between these two runs. Manual interactive shimming of the B0 field was performed for all participants. The system voltage was set at 250 V for all sessions, and the fat suppression was decreased to ensure that the specific absorption rate did not surpass 62% for all functional runs. High-resolution MP2RAGE anatomical images were obtained in the middle of the session with the following parameters: resolution = 0.65 mm isotropic, TR = 5000 ms, TE = 2.51 ms, TI1/TI2 = 900/2750 ms, flip angles = 5/3, iPAT = 2, bandwidth = 250 Hz/Px, echo spacing = 7ms.

#### Data preprocessing

The raw functional data underwent distortion correction using FSL topup (https://fsl.fmrib.ox.ac.uk/fsl/fslwiki/topup). This distortion-corrected data was further processed using the SPM12 toolbox (https://www.fil.ion.ucl.ac.uk/spm/software/spm12). The functional images were realigned, slice-time corrected, co-registered with anatomical images, segmented, and finally normalized to the MNI305 anatomical template. All SPM parameters were set to default and the voxel size after normalization was set to 1.2×1.2×1.2 mm. Before normalization, the data were denoised using GLMdenoise (60), which is known to improve signal-to-noise ratio.

The preprocessed data were concatenated across all runs and modeled using a generalized linear model (GLM) in SPM using the default parameters. For the event-related run, the t-values were estimated for each condition and were used for further analysis instead of beta values.

#### ROI definitions

The regions of interest along the ventral visual pathway were identified by contrasting the activation between conditions from the localizer run. Early visual areas were identified by contrasting scrambled objects with fixation cross. This region was further parsed into V1-V3, and V4 using the anatomical mask from the SPM anatomy toolbox (61). Higher visual area (LOC) was identified as a region that responded more to objects than scrambled objects. The voxels in the LOC region were restricted to the Inferior Temporal Gyrus, Inferior Occipital Gyrus, and Middle Occipital Gyrus. These anatomical regions were obtained from Tissue Probability Map (TPM) labels in SPM 12. VWFA was identified as a region in the left occipital temporal sulcus that responded more to words than scrambled words. For each contrast, a voxel-level threshold of p < 0.001 (uncorrected) was used to obtain a contiguous region.

### MEG data

#### Participants

A total of 18 adult subjects (aged b/w 18-40 years) who were native French speakers participated in this study. They were right-handed and had normal vision with no history of any psychiatric disorders or reading difficulties. Participants provided written informed consent for the MEG study and received monetary compensation. One subject was excluded from this study due to excessive head movements.

#### Stimuli

The stimuli were identical to the fMRI study

#### Task design

As in the fMRI, a series of letter string stimuli were presented to the participants, who had to respond with a button press if all the letters were identical. Each stimulus was shown for 200ms followed by a 300ms blank screen with a fixation cross. In a run, each stimulus was repeated 15 times, plus 60 target trials. The study comprised three such runs in total.

#### Data acquisition

Participants performed the tasks while sitting inside an electromagnetically shielded room. The magnetic component of their brain activity was recorded with a 306-channel, whole-head MEG by Elekta Neuromag (Helsinki, Finland). 102 triplets, each comprising one magnetometer and two orthogonal planar gradiometers composed MEG helmet. The brain signals were acquired at a sampling rate of 1000 Hz with a hardware highpass filter at 0.03 Hz. Eye movements and heartbeats were monitored with vertical and horizontal electrooculograms (EOGs) and electrocardiograms (ECGs). Head position inside the helmet was measured at the beginning of each run with an isotrack Polhemus Inc. system from the location of four coils placed over frontal and mastoidian skull areas.

#### Data preprocessing

The MEG data underwent several preprocessing steps. Initially, the data was inspected to identify any sensors showing excessive noise or poor signal quality. This was performed using a custom script in which sensors displaying deviations from the median variance across time by more than 6 standard deviations were identified as deviant sensors and excluded from further analysis.

To remove environmental noise and artifacts related to head movements, Maxwell filtering was applied using the MaxFilter tool in MNE-Python. This step compensated for head position changes during the recording session by using the continuous head position indicator (cHPI) coil signals. The data were processed with signal space separation (SSS) to suppress external interference and enhance the signal-to-noise ratio. A finite impulse response bandpass filter was then applied to the data to focus on the frequency range of interest (1-40 Hz) and eliminate unwanted noise.

Signal space projection (SSP) techniques were employed to further improve data quality and reduce artifacts. SSP vectors were computed by identifying signal components, such as eye blinks or heartbeats. These SSP vectors were then projected out from the MEG data to remove the corresponding artifacts.

#### Eye tracking

During the MEG acquisition, participants were instructed to fixate on the central cross while the items were flashed. Their gaze was monitored online using EyeLink 1000 eye-tracker device (SR research). Eye-tracking data was collected for 16 out of 18 participants

#### Decoding analysis

The stimuli used in fMRI and MEG experiments varied along the following factors: absolute word position, absolute letter position, ordinal letter position, and letter identity. To analyze the neural signals corresponding to these factors, a linear classifier was used. Specifically, logistic regression with a “liblinear” solver was performed after scaling the data using “robustscalar” from the sci-kit learn package in Python. To avoid overfitting, a 3-fold cross-validation was performed using the MNE function “cross_val_multiscore”. For decoding the position information, the classifier was trained on the data from any two letter pairs, and tested on the third one. Similarly, for decoding letter identity, the classifier was trained on any two ordinal positions and tested on the remaining one. In fMRI, the voxels within each region were used as features, and in MEG, the channels were used as features with data averaged within 10ms time-bins.

#### Estimating neural dissimilarity

In fMRI, for a given ROI and subject, pair-wise dissimilarity between a stimulus pair was computed using correlation distance metric (i.e., 1 – r); where r is the correlation coefficient between the activity pattern across voxels. Similarily, in MEG, the signal across each channel was first baseline corrected using MNE function “mne.baseline.rescale” by subtracting the mean and dividing by the standard deviation of the baseline values (zscore). The correlation distance was then estimated using channels as features

