## Supplementary data for "Dissecting the cortical stages of invariant word recognition"

**Dissecting the neuronal mechanisms of invariant word recognition**

**Supplementary materials**

- Supplementary Figure 1: Stimuli used in Brain imaging studies
- Supplementary Figure 2: Behavioral Task Performance
- Supplementary Figure 3: Representational similarity space predicted by literate CNN
- Supplementary Figure 4: MEG model fits for each sensor
- Supplementary Figure 5: Searchlight analysis between CNN and fMRI

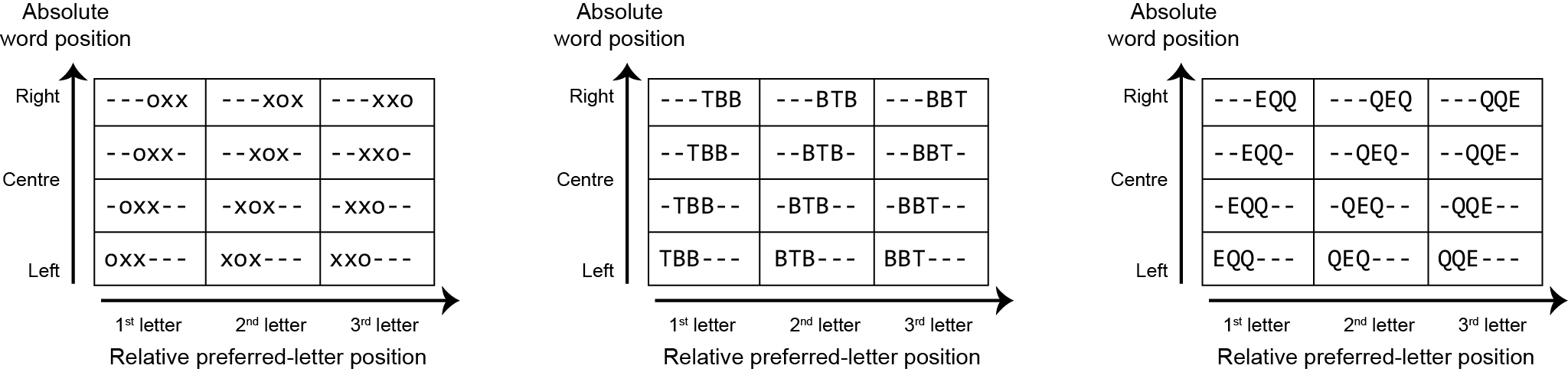

**Figure S1: Stimuli used in brain imaging studies.** Schematic of the stimuli used to dissociate position coding. The word position varies across rows, and the preferred letter position within a string varies across columns. ‘-’ represents blank space. Overall, there are 36 unique stimuli.

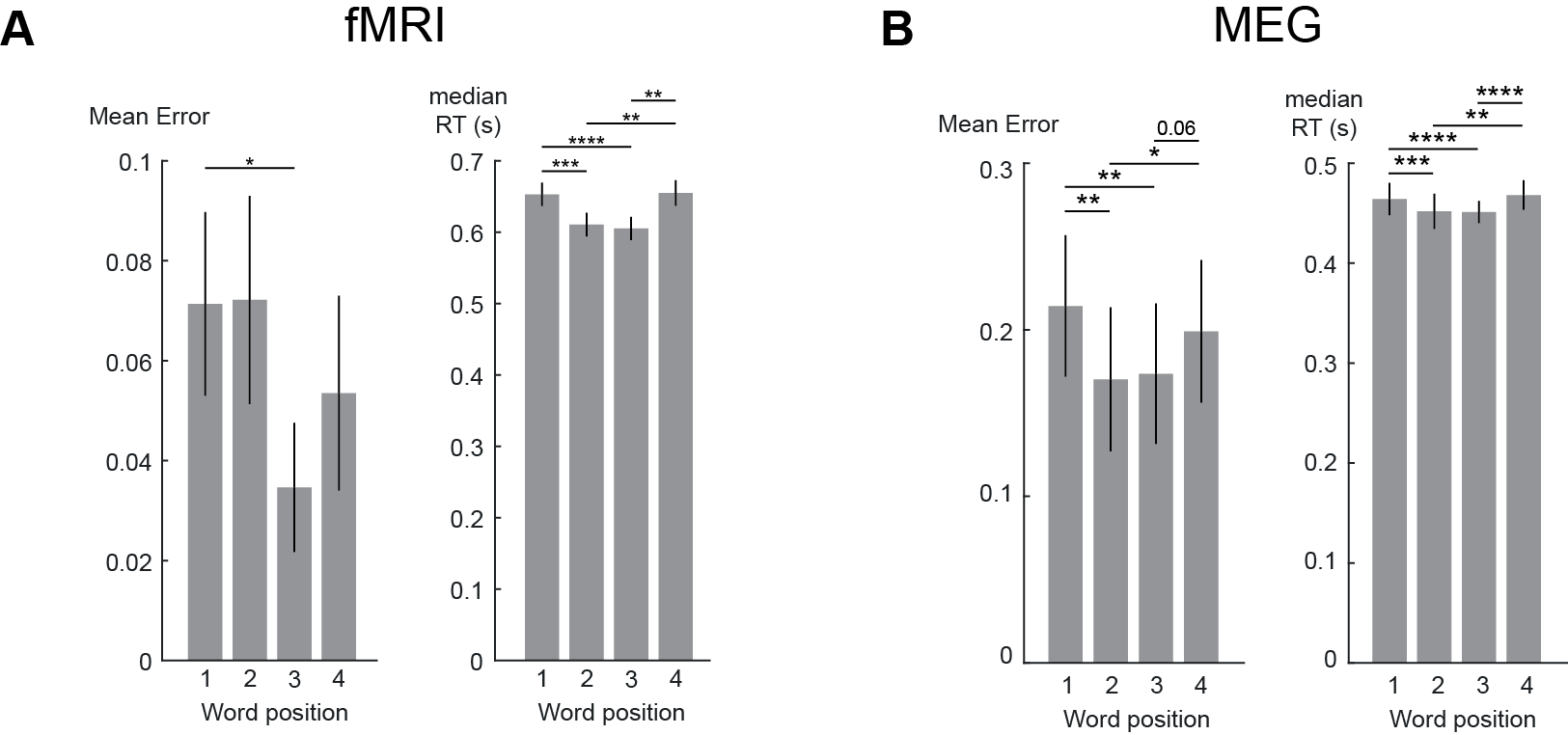

**Figure S2: Behavioral task Performance.**

(A) During fMRI data acquisition, subjects were instructed to respond with a button press when all the letters in a stimulus were identical. The stimuli could appear at any word position during task trials. Subjects exhibited low error (left) and faster response times (right) when the stimuli were presented at the fovea. Each run included 10% task trials. Error bars indicate standard deviation across subjects and the asterisks indicate statistical significance using paired t-test (* P < 0.05, ** p < 0.005, *** p < 0.0005, etc).
(B) Same as (A) but for MEG data. Faster stimulus presentation with an inter-trial interval of 300 ms resulted in a speed-accuracy trade-off: in MEG, compared to fMRI, subjects were less accurate but responded faster.

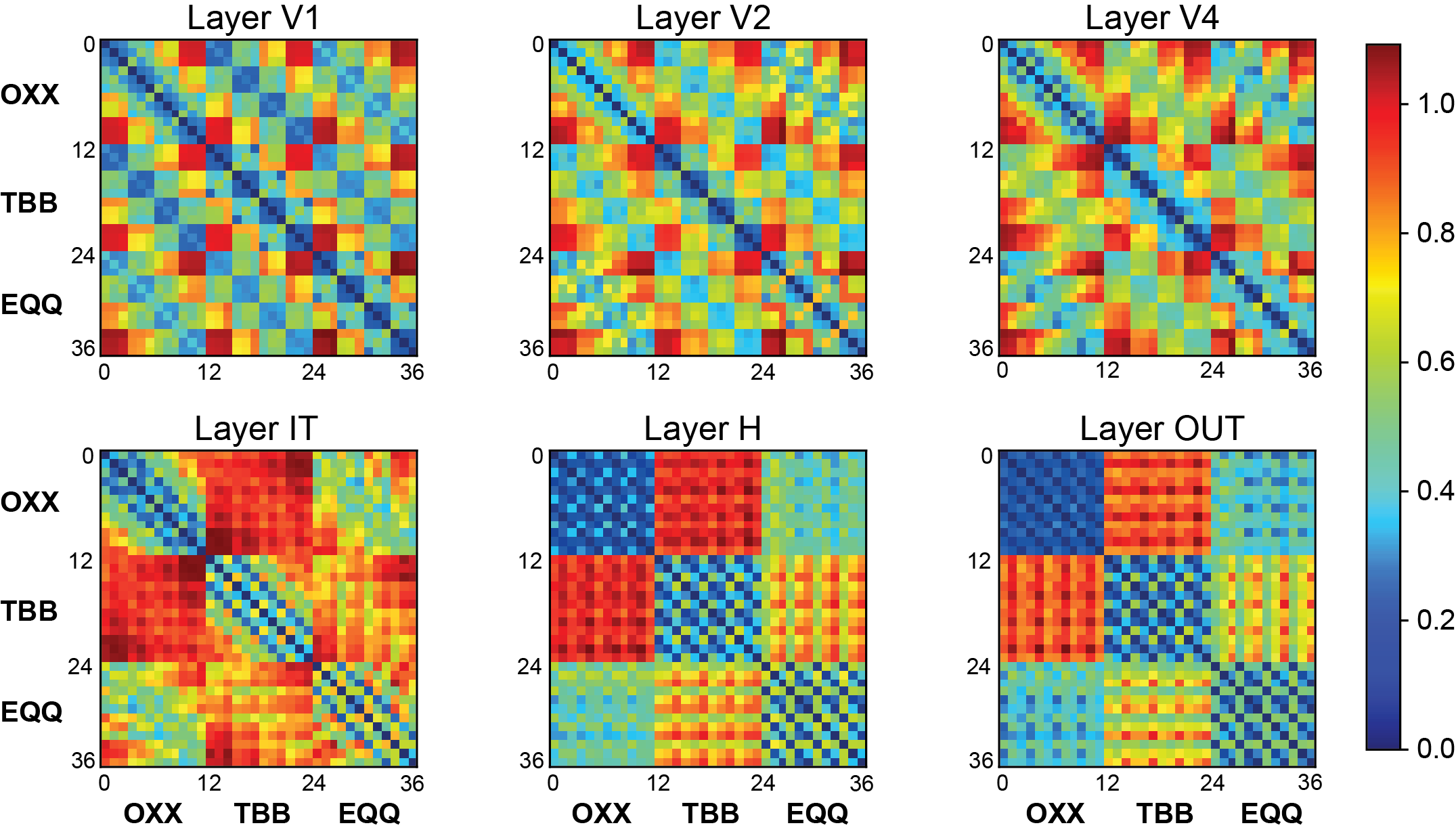

**Figure S3: Representational similarity space predicted by literate CNN.** Representation Dissimilarity Matrices (RDMs) predicted by different layers of the literate network across the 36 stimuli that were used in the brain-imaging experiments. The correlation distance metric (1-r) is plotted according to the color scale at right. In the early layers, word position and absolute letter positions are encoded irrespective of the specific letter identities. The IT layer demonstrates partial invariance to word position and encodes letter identity. Complete position invariance is observed from the H layer.

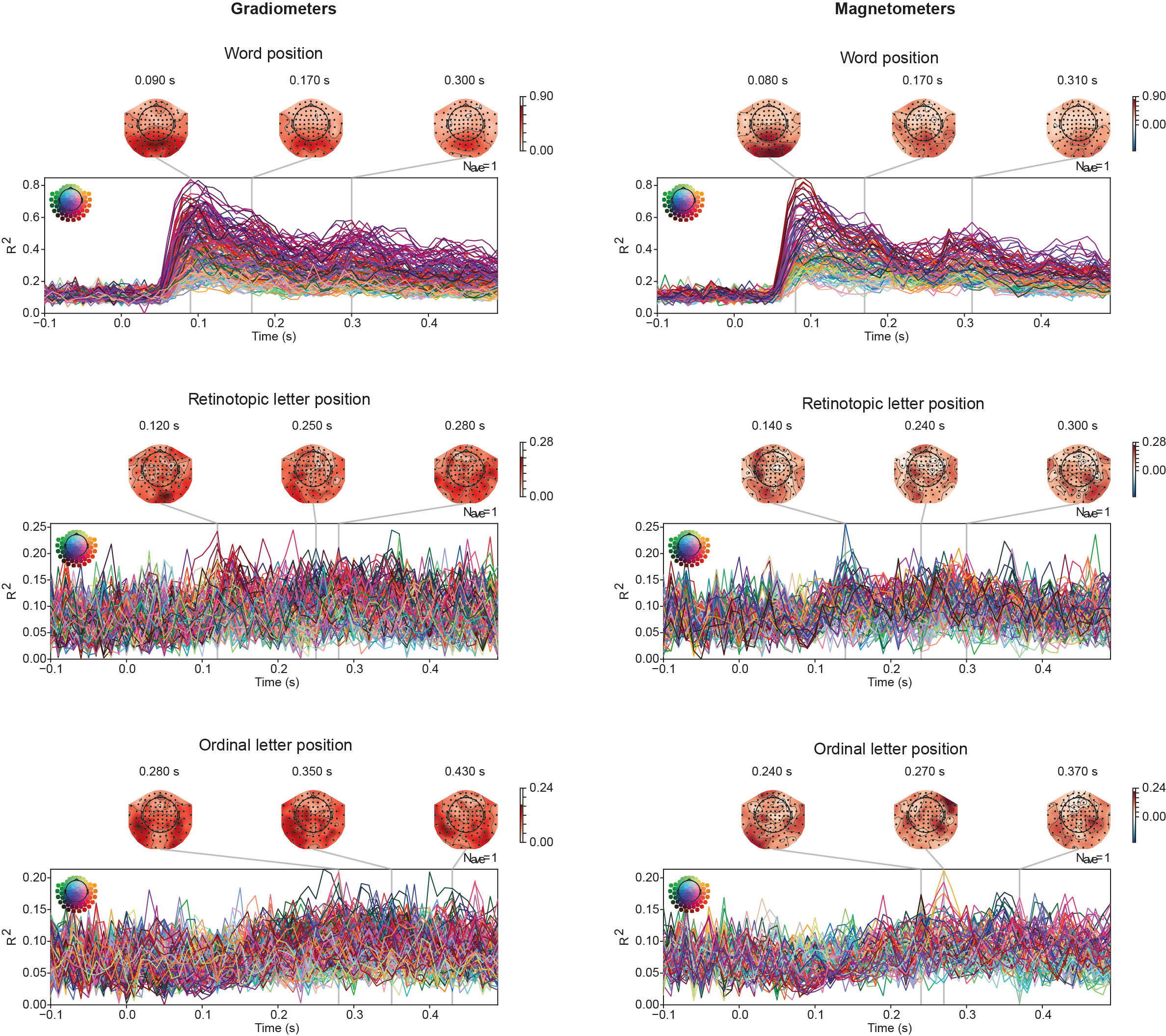

**Figure S4: MEG model fits for each sensor.** Temporal evolution of encoding model fits from MEG recordings, showing partial variance (R²) from -100 to 500 ms relative to stimulus onset for each of the Gradiometer (left) and Magnetometer (right) sensors. We observe that the magnitude of model fits for word position is highest in sensors located at the back of the topomap, corresponding to early visual areas. Conversely, the sensors with the highest magnitude of model fits for ordinal position are located slightly laterally, corresponding to anterior temporal regions

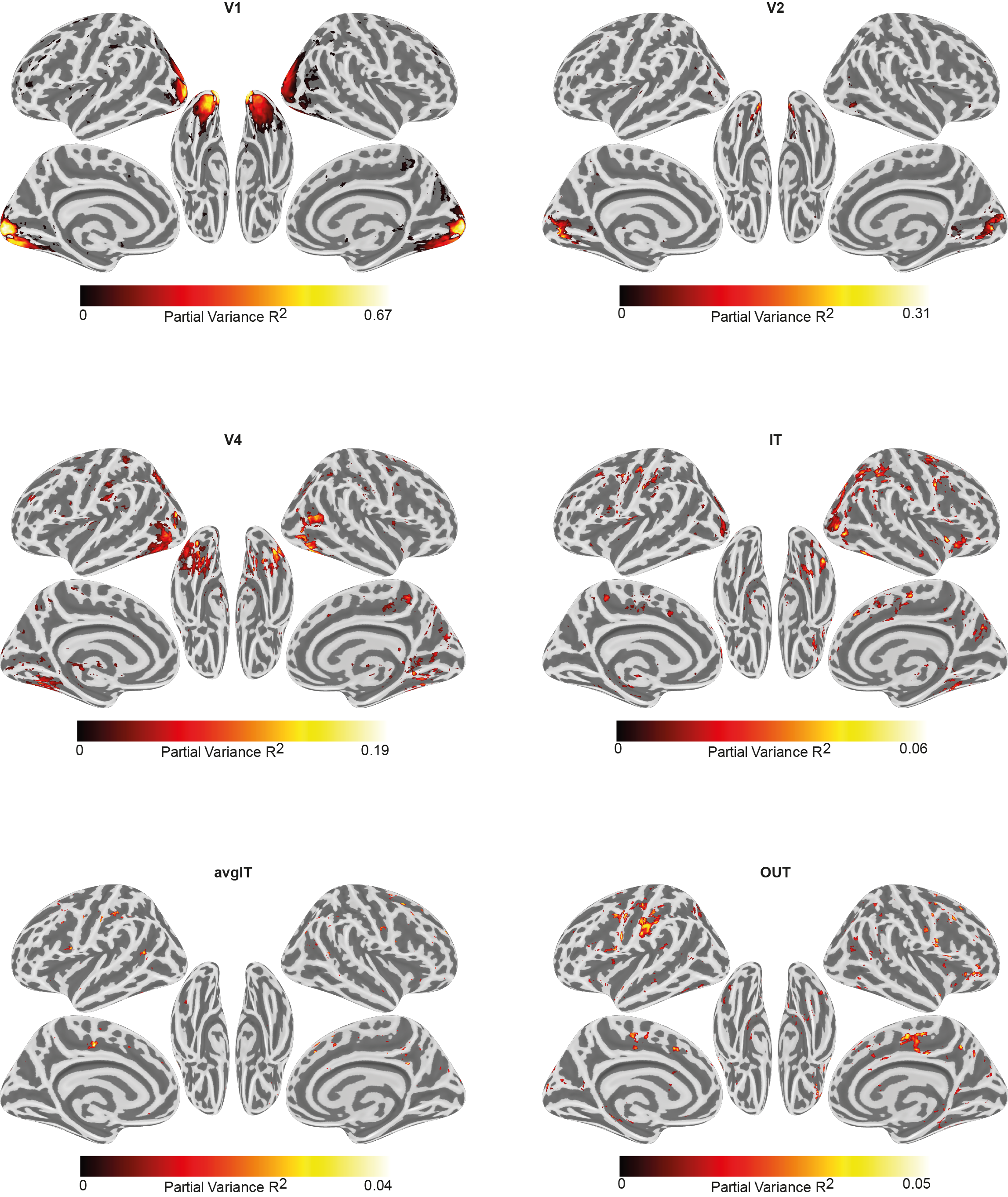

**Figure S5: Searchlight analysis between CNN and fMRI.** Searchlight maps of the partial variance estimated via hierarchical regression between the representation space of CNN layers and neural activity at each voxel. These maps are thresholded to indicate the significant clusters after FDR correction (p<0.05)
